# Genome Editing Outcomes Reveal Mycobacterial NucS Participates in a Short-Patch Repair of DNA Mismatches

**DOI:** 10.1101/2023.10.23.563644

**Authors:** Tanjina Islam, Eric A. Josephs

## Abstract

In the canonical DNA mismatch repair (MMR) mechanism in bacteria, if during replication a nucleotide is incorrectly mis-paired with the template strand, the resulting repair of this mis-pair can result in the degradation and re-synthesis of hundreds or thousands of nucleotides on the newly-replicated strand (long-patch repair). While mycobacteria, which include important pathogens such as *Mycobacterium tuberculosis*, lack the otherwise highly-conserved enzymes required for the canonical MMR reaction, it was found that disruption of a mycobacterial mismatch-sensitive endonuclease NucS results in a hyper-mutative phenotype, which has led to the idea that NucS might be involved in a cryptic, independently-evolved DNA MMR mechanism. It has been proposed that nuclease activity at a mismatch might result in correction by homologous recombination (HR) with a sister chromatid. Using oligonucleotide recombination, which allows us to introduce mismatches during replication specifically into the genomes of a model for *M. tuberculosis, Mycobacterium smegmatis*, we find that NucS participates in a direct repair of DNA mismatches where the patch of excised nucleotides is largely confined to within ∼5 - 6 bp of the mis-paired nucleotides, which is inconsistent with mechanistic models of canonical mycobacterial HR or other double-strand break (DSB) repair reactions. The results presented provide evidence of a novel NucS-associated mycobacterial MMR mechanism occurring *in vivo* to regulate genetic mutations in mycobacteria.

## INTRODUCTION

In 2021, there were 10 million incident cases of tuberculosis (TB) worldwide, about 3.9% (450,000) of which were resistant to first line drug rifampicin or multiple antibiotics; this figure represents about 18% of all previously-treated cases (1). *Mycobacterium tuberculosis*, the pathogenic bacteria that causes TB, acquires drug resistance exclusively through chromosomal mutation events, especially single-nucleotide polymorphisms (SNPs) (2,3). However, there is currently an incomplete understanding of the molecular processes that govern the mechanisms of mutation and mutation avoidance in mycobacteria. In nearly all other organisms, except for some archaea and actinobacteria which includes mycobacteria, the rate of genetic mutation is tightly controlled by a MutS/MutL-coordinated DNA mismatch repair (MMR) mechanism (4). Immediately after replication, the MMR reaction corrects any incorrectly incorporated or ‘mismatched’ nucleotides that would become permanent genetic mutations if left unrepaired. For example, in the well-studied DNA MMR reaction in *Escherichia coli*, after MutS recognizes a mismatched nucleotide in double-stranded DNA, MutS together with MutL activates latent nicking endonuclease MutH, which then nicks hemi-methylated DNA at d(GATC) sites on the unmethylated strand—an epigenetic signal that discriminates the newly-replicated from the template strand (5). Helicases and exonucleases are loaded at the nicked site and degrade the newly-replicated strand through the mismatch so the strand can be re-synthesized along that tract. Strand degradation and resynthesis can be coordinated between the epigenetic signals and the mismatch even if they are separated by hundreds or thousands of nucleotides in what is known as “long-patch” repair. During senescence, MMR proteins also contribute to the recognition and repair of chemically damaged nucleotides and inhibit improper recombination events between divergent sequences (6-8).

In cells with inactivated MMR pathways, the basal mutation rate is increased100-fold relative to cells with active MMR (5). However, most actinobacteria have similar basal mutation rates as bacteria with MutS/MutL-coordinated MMR pathways, despite lacking any known homologues of MMR proteins MutS or MutL (9-11). This was originally thought to be a result of unusually high-fidelity mycobacterial DNA polymerases (12). Rather, it was recently discovered (13) that disruption of the mycobacterial gene *nucS*, which encodes an endonuclease sensitive to mismatched nucleotides, resulted in the same hyper-mutative and hyper-recombinative phenotypes observed for other bacteria in which *mutS* or *mutL* had been disrupted. This finding lead to the hypothesis that NucS might be an involved in an evolutionarily-divergent DNA MMR mechanism. *In vitro*, actinobacterial NucS enzymes recognize and cleave double-stranded DNA containing mis-paired dG-dG, dG-dT, and dT-dT nucleotides, and endonuclease activity requires association with the beta clamp of the replisome (14,15). However, not much is known about the molecular mechanisms by which NucS activity results in the suppression of a hyper-mutative phenotype. It has been proposed that double-strand breaks (DSBs) caused by NucS when nucleotides are mismatched during replication could be repaired by homologous recombination (HR) with a sister chromatid through gene conversion (16). NucS is evolutionarily related to an archaeal mismatch-sensitive endonuclease EndoMS which has also been bioinformatically associated with the enzymes responsible for HR in archaea (17). Alternatively, in principle, a mismatch-sensitive endonuclease might simply induce cell death in members of the bacterial population that are hyper-mutative for other reasons, rather than participating in the correction and repair of the mismatched nucleotides itself.

Typically, DNA MMR activity and hypermutation are characterized using spontaneous antibiotic resistance assays that can provide an estimate of mutation rates under certain conditions by observing the frequency at which a microorganism acquires one of a spectrum of mutations that result in resistance to certain antibiotics, such as mutations in the gene for the beta subunit of RNA polymerase *rpoB* that result in resistance to rifampicin (13). Mutation accumulation studies using whole genome sequencing have also been performed to understand the role NucS plays in genome maintenance and mutation, which confirmed the role of NucS in limiting transition mutations in *Mycobacterium smegmatis,* a non-pathogenic model of *M. tuberculosis,* that would be caused by dG-dG, dG-dT, and dT-dT mis-pairing during replication (18). Using a genome engineering technique known as oligonucleotide recombination (OR) (Figure 1A) (19-22), where engineered oligonucleotides that contain mismatched nucleotides with the lagging-strand template are introduced into bacteria and incorporated into the genome during replication, we are able to directly evaluate different molecular mechanisms associated with NucS-associated mutation avoidance in *M. smegmatis*. We find that genomic mismatches introduced during OR are rejected in a manner consistent with the biochemical association of NucS with various combinations of mismatched nucleotides (14,15), but that these mismatches can be repaired in a way that is inconsistent with both the long-patch repair of canonical MMR and canonical mycobacterial DSB repair mechanisms like HR or non-homologous end-joining (NHEJ) (23,24). Our results show that a mycobacterial MMR can occur involving NucS within a short patch (less than about ∼6 - 9 nt) of the mis-paired nucleotides. We also find that that compound mismatches (25) introduced *via* OR can help to evade NucS-mediated repair and allow for genome editing of mycobacterial species in a manner that we expect will prove useful for probing mycobacterial biology and virulence. The results presented here provide direct evidence that NucS can participate in a mycobacterial MMR reaction that from the outcomes of OR gene editing appears to be mechanistically distinct from repair by HR or other canonical mycobacterial DSB repair mechanisms.

**Figure 1.**
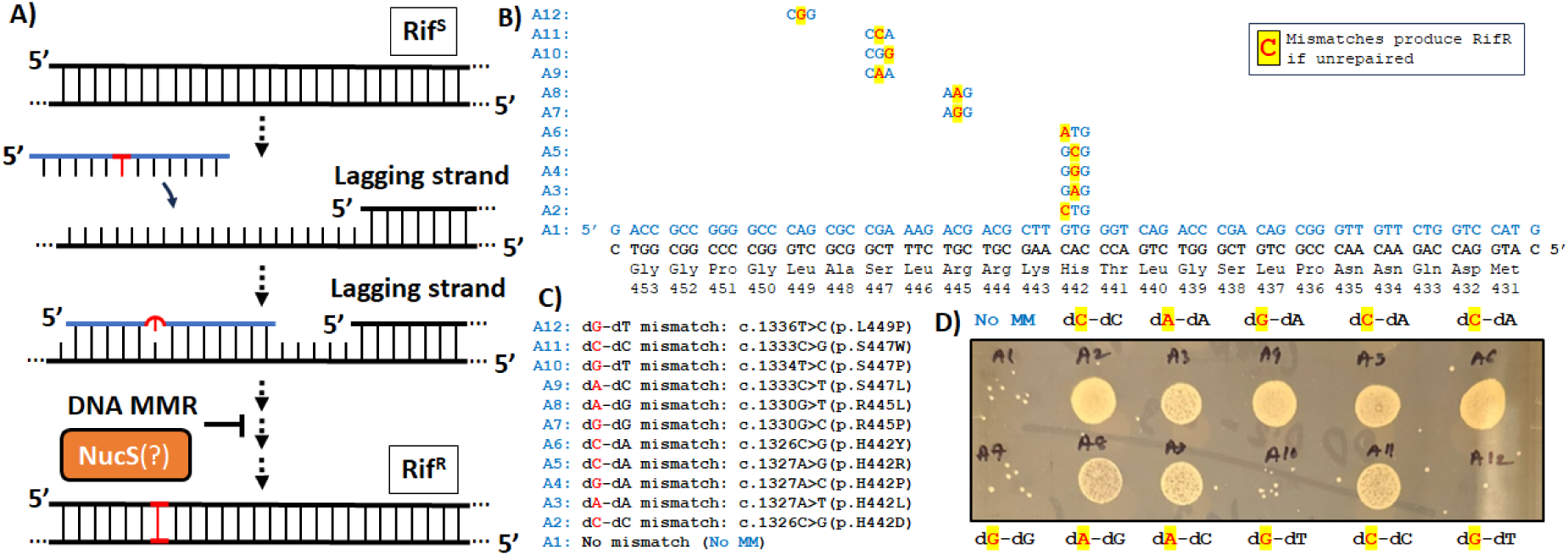
*Oligonucleotide recombination (OR) to probe the involvement of NucS in the repair of mismatched nucleotides in M. smegmatis. A) A schematic model of OR, which is attenuated by DNA mismatch repair (MMR) processes. Mismatches left unrepaired after introduction into the rpoB gene by OR results in a transition from a rifampicin-sensitive (Rif^S^) phenotype to a rifampicin resistant (Rif^R^) phenotype. B) 12 oligonucleotides (blue) differ by a single nucleotide to introduce different varieties of mismatched nucleotides that produce different known Rif^R^ mutations. Only the sequence differences with the control oligonucleotide (A1) which base-pairs perfectly with the lagging-strand template and surrounding codon are shown for A2-A12. Below, lagging strand template sequence (black) with codons and numbering for M. smegmatis rpoB. C) Different known Rif^R^ mutations introduced by the oligonucleotides. D) M. smegmatis with Rif^R^ phenotype are efficiently generated when using oligonucleotides that introduce dC-dC, dA-dA, dG-dA, dC-dA, dA-dC, and dA-dG mismatches, but not dG-dT or dG-dG mismatches, known targets of NucS*.

## RESULTS

During OR (22,26,27), short oligonucleotides are introduced into cells and base-pair with the lagging-strand template, and if there are any mismatched nucleotides on the oligonucleotide they become permanent genetic mutations if left unrepaired before a subsequent round of replication (Figure 1A) (19-21). While genome engineering by OR has been demonstrated in *M. tuberculosis* and *M. smegmatis* (28-31), OR is not commonly used in mycobacteria, perhaps because of an incomplete understanding of mismatch repair processes in mycobacteria (32). There is significant evidence that OR occurs during replication in mycobacteria and other bacteria, as there is a strong bias in OR efficiency favouring oligonucleotides that base-pair with the lagging-strand template *vs.* the leading-strand template (28), and OR does not occur when the complementary DNA is not actively replicating (20). A re-evaluation of previously-reported OR efficiencies in mycobacteria in light of the discovery of NucS and its biochemical characterization that it cleaves dT-dT, dG-dT, and dG-dG mismatches in double-stranded DNA suggested that OR might be attenuated by the activity of NucS (not shown) (28).

Because OR has been used to probe cellular processes in *E. coli*, yeast, and human cells (25,33-38)—in particular, DNA MMR processes (39,40)—we sought to determine whether oligonucleotides with mismatches introduced during OR into *M. smegmatis* were rejected by NucS. We designed a series of 11 nearly-identical oligonucleotides that contained various mis-pairing nucleotides across 6 codons in a region of *M. smegmatis* gene *rpoB* to generate mutations that are known to result in resistance to the antibiotic rifampicin (41,42)if the mismatched nucleotides are left unrepaired (Figure 1B-C).

Log-phase *M. smegmatis* were individually electroporated with these eleven oligonucleotides a control oligonucleotide containing no mis-pairing nucleotides and therefore designed to cause no mutation, allowed to recover, then plated on solid media containing rifampicin (28). We saw a significant enhancement in the population of survivors compared to the control across oligonucleotides that introduced dC-dC, dA-dA, dG-dA, dA-dG, dC-dA, and dA-dC mismatches (Figures 1D and S1). Each of those mismatched nucleotide pairs are not recognized by NucS *in vitro* (14,15). This enhancement in the number of rifampicin-resistant survivors was not observed in oligonucleotides that introduced dG-dG or dG-dT mismatches that, *in vitro,* are cleaved by NucS. This provided initial evidence that, as expected, mismatches introduced using OR are rejected through a NucS-mediated mechanism, which was further supported by additional evidence provided below.

We then performed OR using oligonucleotides that introduced both a (NucS-inactive) dA-dC mismatch to produce rifampicin resistance if unrepaired and a nearby (NucS-active) dG-dT mismatch that would generate a synonymous mutation in *rpoB* if unrepaired (Figure 2A). Based on the model that a DSB caused by NucS at a mismatch would be repaired by HR (16), we expected that these oligonucleotides would be less than half as effective as those introducing a dA-dC mismatch alone as the dA-dC mismatch might be repaired “collaterally” by gene conversion or end resection at the initiation of HR (23). However, *M. smegmatis* that had oligonucleotides to introduce dG-dT mismatches as close as 5 or 6 bp away from the dA-dC mismatch also showed significant levels of rifampicin resistance, similar to oligonucleotides that introduced dA-dC mismatches alone (Figures 2B-C and S2). Sanger sequencing of the population (kept in liquid media containing rifampicin without plating) confirmed that the rifampicin resistance was largely a result of the specific *rpoB* c.1327A>G (p.H442R) mutation introduced *via* the dA-dC mismatch, while there was no evidence of the other dA-to-dG mutations that would have been introduced by the dG-dT mismatch on the same oligonucleotide (Figure 2D). In contrast, if an oligonucleotide that would introduce two NucS-inactive mismatches (dA-dC for rifampicin resistance and dA-dG for a synonymous mutation in *rpoB*), the levels of mutation at both those sites were correlated.

**Figure 2.**
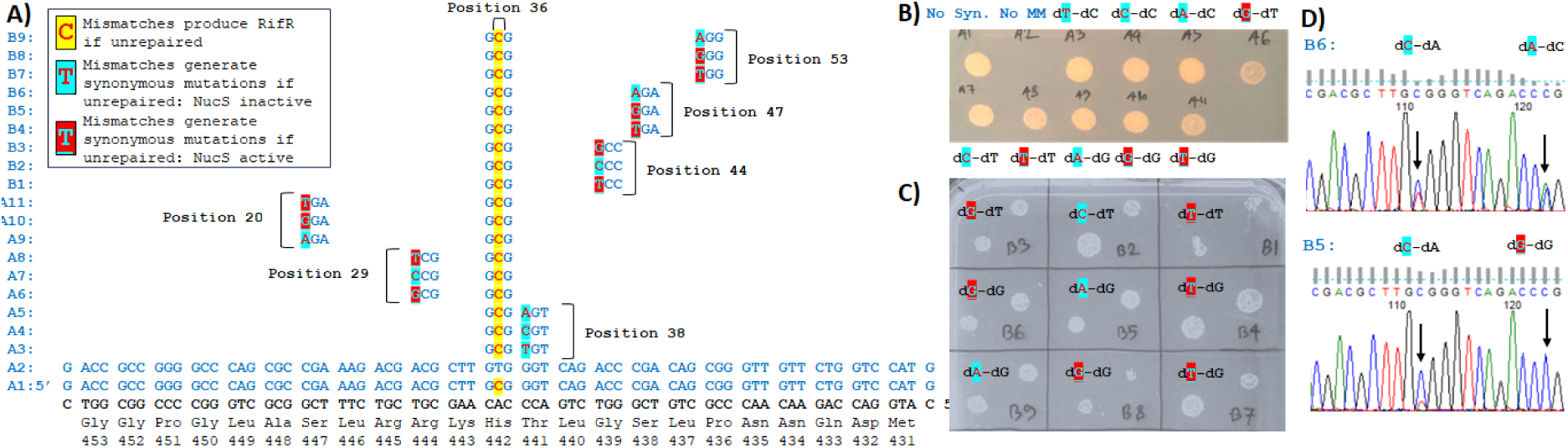
*Oligonucleotides which introduce NucS-active mis-pairs near a mis-pair to generate a rifampicin-resistance mutation do not affect OR. A) Oligonucleotides (blue, see Figure 1 caption) where NucS-active (red with blue letters) and NucS-inactive (blue with red letters) mis-pairs are introduced to generate synonymous mutations in rpoB along with mis-pairs to generate rifampicin resistance (red). Positions labelled relative to oligonucleotide 5’-. B and C) The molecular identify of the synonymous mutation (highlighted nearby) does not affect the efficiency of acquiring rifampicin resistance from these oligonucleotides after recovery and plating on solid media with rifampcin. D) From Sanger sequencing of M. smegmatis recovered together in liquid media with rifampicin (not isolated as colonies), synonymous mutations caused by NucS-inactive mismatches are correlated with rifampicin-resistance mutation caused by a NucS-inactive mismatch, but synonymous mutations caused my NucS-active mismatches are not*.

While successful OR itself occurs with low frequency, the above result meant that we could select for the rifampicin resistance mutation caused by a NucS-inactive dA-dC mismatch to isolate and enrich cells with successful oligonucleotide incorporation. This would allow us to quantify the presence or absence of mutations nearby that would also be introduced on the same oligonucleotide and significantly increase sequencing depth for next-generation sequencing so that we could perform experiments using multiple oligonucleotides with higher throughput, and so that the mutational results from different variations of oligonucleotides could be compared within the same experiment. It also allows us to perform these types of experiments and compare with strains with *nucS* gene knocked out (43), as the success rate of OR approaches the rate of spontaneous rifampicin resistance in these strains.

Repeating this experiment using pooled oligonucleotides (from Figure 2A) that would introduce a rifampicin-selective dA-dC mismatch flanked at various positions by different synonymous mismatches (Figure 3) and comparing the results between wild-type and a *nucS*-knockout stain of *M. smegmatis* confirmed that only the dG-dG, dG-dT, and dT-dT mismatches on those oligonucleotides were removed in a NucS-specific manner. The results were highly reproducible and revealed a slight but detectable defect in the repair of dG-dT mismatches relative to dG-dG and dT-dT mismatches, the mutations caused by which were essentially undetected, and of a slight difference between dT-dG (dT base-paired with template dG) and dG-dT (dG base-paired with template dT). These results also confirmed the measured biochemical activity of NucS matched the repair of mismatches by *M. smegmatis in vivo* (14,15). Therefore, we conclude that NucS-active mismatches introduced during OR are being specifically removed by a NucS-mediated mechanism. Surprisingly, there were also no orientation effects (locating NucS-active mismatches either 5’- to 3’- relative the dA-dC) as one might expect during repair by HR (*i.e.*, see Figure 7): based on the cleavage pattern of NucS (nicking the DNA two nucleotides 5’- the mismatch on both strands) (14,15,17), depending on whether or not the NucS-active mismatch was 5’- or 3’- of the other one might expect NucS-active mismatches to be repaired during single crossover HR on either different fragments or on the same fragment. In these and subsequent experiments, we also found no evidence of the hallmarks of other major double-strand break (DSB) repair mechanism by non-homologous end-joining (NHEJ), that is, no nucleotide insertions or deletions at the site of the repaired mismatches (Figure S3) (44).

**Figure 3.**
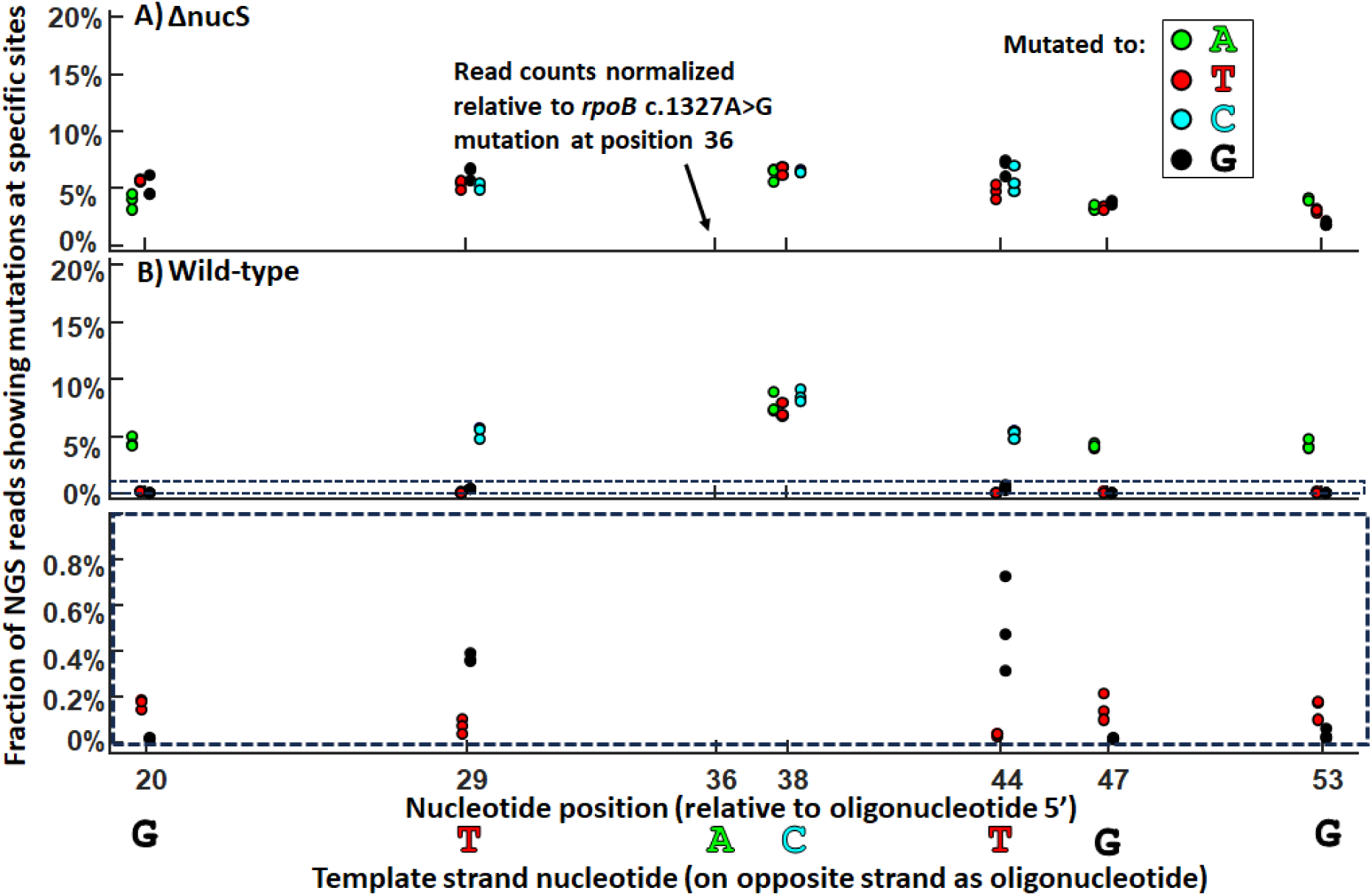
*Next-generation sequencing (NGS) results after pooling the oligonucleotides in Figure 2A for OR and rifampicin selection. Note that the results presented show 3 biological replicates (if fewer than three dots are observed, it is because they are overlapping). A) In a M. smegmatis strain with nucS gene deleted, each mutation caused by all of the oligonucleotides are represented at approximately equal frequencies. B) In the wild-type strain, frequency of mutations that would occur as a result of dG-dG, dT-dG, and dT-dT mismatches sharply decreases. Below: highlighting low frequency mutations: it appears dT-dT = dG-dT < dT-dG < dG-dT with regards to repair efficiency by NucS*.

We then performed OR using oligonucleotides that would introduce (Figures 4A and S4):

i. a dA-dC mismatch that should introduce a rifampicin resistant phenotype if unrepaired;
ii. a NucS-active dT-dG mismatch located either 5’- or 3’- of (i) that would produce synonymous mutation if unrepaired; and
iii. one or more NucS-inactive mismatches (e.g., dA-dC, dC-dC, dA-dA, dT-dC) that would produce synonymous mutations in *rpoB* if unrepaired, at various positions relative to (i) and (ii).

**Figure 4.**
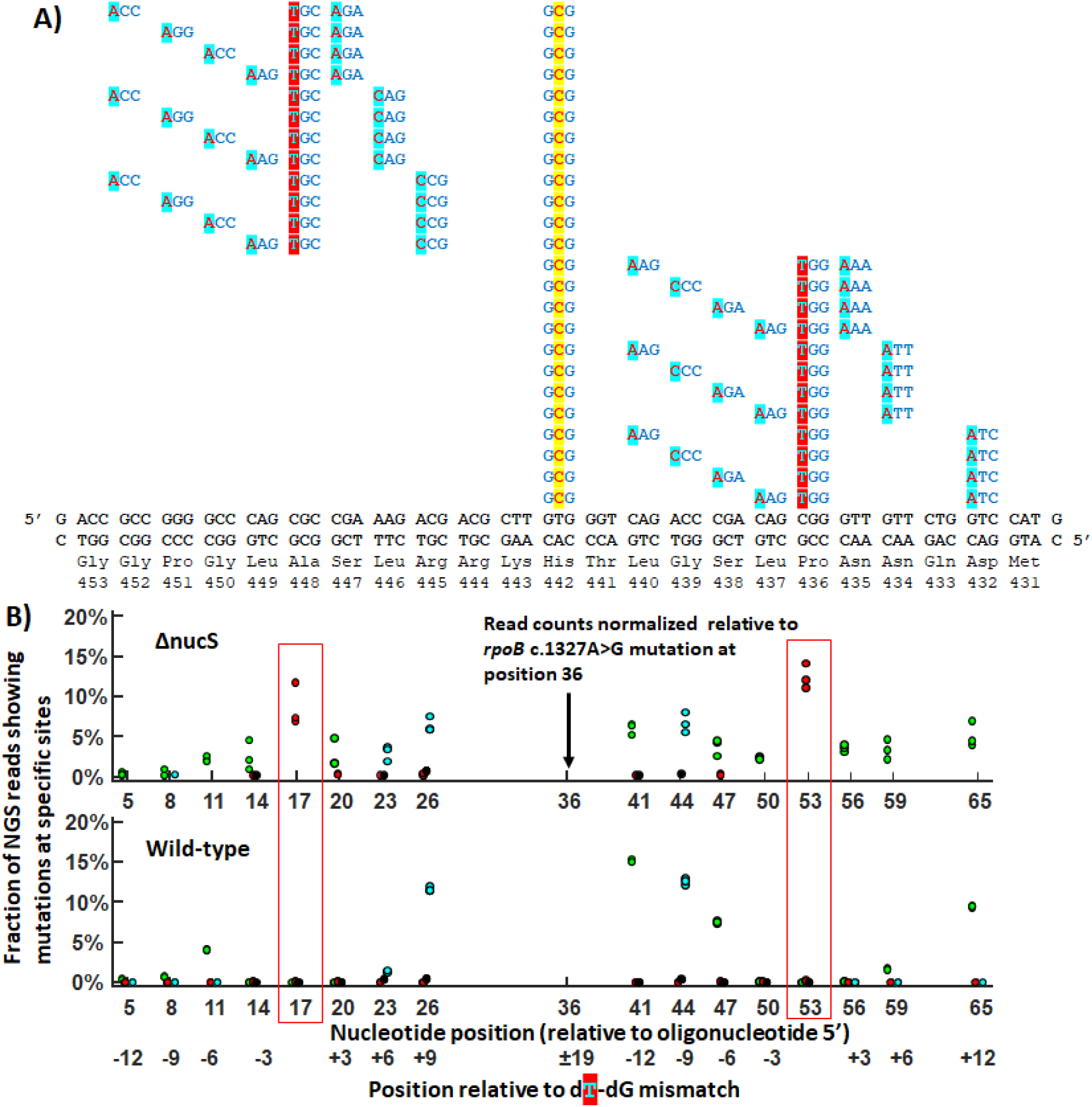
*NucS-associated MMR collaterally repairs NucS-inactive mismatches within 6 nucleotides of a NucS-active mismatch but not outside a 6 – 9 nt region of the NucS-active mismatch. A) Pooled oligonucleotides (blue, see Figure 1 caption) that contain i) a dA-dC mismatch that should introduce a rifampicin resistant phenotype if unrepaired; ii) a NucS-active dT-dG mismatch located either 5’- or 3’- of (i) that would produce synonymous mutation if unrepaired; and iii) two NucS-inactive mismatches (e.g., dA-dC, dC-dC, dA-dA, dT-dC) that would produce synonymous mutations in rpoB if unrepaired, at various positions relative to (i) and (ii). B) Mutations generated by both NucS-active (boxed in red) and NucS-inactive mismatches within 3 nt of a NucS-active mismatch are significantly depleted in the NucS-active strain. Mutations generated by NucS-inactive mismatches > 6 nt away are largely unaffected, though there is a slight effect 9 nt 3’- of the NucS-active mis-pair. Note that the results presented show 3 biological replicates (if fewer than three dots are observed, it is because they are overlapping). Figure S4 has a variation of this experiment*.

After selection in rifampicin and next-generation sequencing (NGS), we report several unexpected findings. As before, in a NucS-dependent manner we found that the dG-dT mismatches were efficiently repaired when flanked by NucS-inactive nucleotides, but that NucS-inactive mismatches were found to be repaired “collaterally” when located 3 nt away from the mismatched dG-dT (Figure 4B and S4). Mutations caused by a dG-dT mis-pair and dA-dG mis-pairs 3 nt away both 5’- and 3’- away from the dG-dT are significantly depleted from sequencing reads in wild-type (NucS active) *M. smegmatis*. This was observed in both cases where the dG-dT mismatch is located either 5’- and 3’- of the *rpoB* c.1327A>G site, and in all oligonucleotides. However, flanking mis-matched nucleotides located 6 nt away or farther from the dG-dT mismatch on the same oligonucleotide were largely not repaired, although there appeared to be and incomplete collateral repair of 3’- but not 5’- NucS-inactive mis-pairs located 6 nt away (but not the one 9 nt away). When looking at the correlation between mutations that are observed in the sequencing reads (Figure 5), we find a significant fraction (∼15%) have mutations both 5’- and 3’- the site of the dG-dT mis-pair— and indeed located as close as 6 nt away from the site of the dG-dT mis-pair—but no mutation caused by the dG-dT mis-pair. This would imply that a NucS-associated MMR process can occur within a notably short patch of DNA.

**Figure 5.**
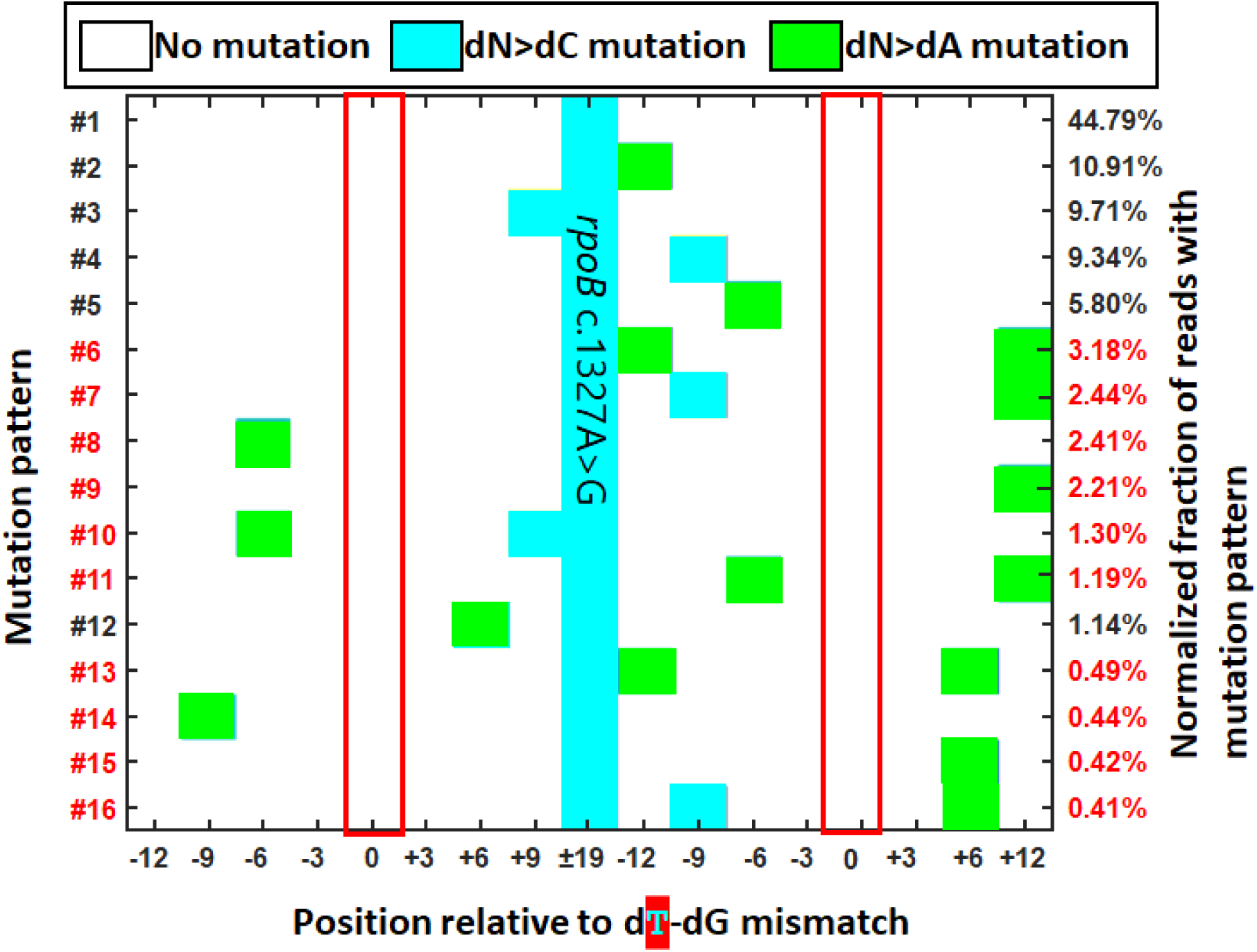
*Mutation patterns from next-generation sequencing (NGS) with relative frequency >0.1%, the average of 3 biological replicates observed after OR with oligonucleotides in Figure 4A. Red boxes highlight the position of the dT-dG mismatches. Patterns marked with red labels have mutations introduced to both sides the site of the dT-dG mismatch, with no associated dC>dT mutation*.

Lastly, in other species, OR can be performed in a way to evade DNA MMR and increase OR efficiency for mis-pairs that would otherwise be very readily repaired before mutations were made permanent (25,36). This is done by exploiting the inability of MutS and MutS homologues to recognize dC-dC mismatches or complex lesions, and it had been previously found that introducing complex mismatches, where multiple dC-dC mismatches flank the mismatch of interest at locations where they would generate synonymous mutations, or a series of multiple consecutive mis-pairs, could be used to increase OR efficiency of mutations caused by MutS-recognized mismatches. Because *M. smegmatis* NucS has a different spectrum of mismatches recognized than MutS, we investigated which types of complex lesions might evade NucS-mediated repair (Figure 6). We found that introducing 3 consecutive or semi-consecutive lesions within 6 nt of each other near a NucS-active mismatch, whether or not they are NucS-active or inactive, appeared to be able to most efficiently evade NucS-mediated repair even when including, dG-dG, dG-dT, dT-dT mismatches.

**Figure 6.**
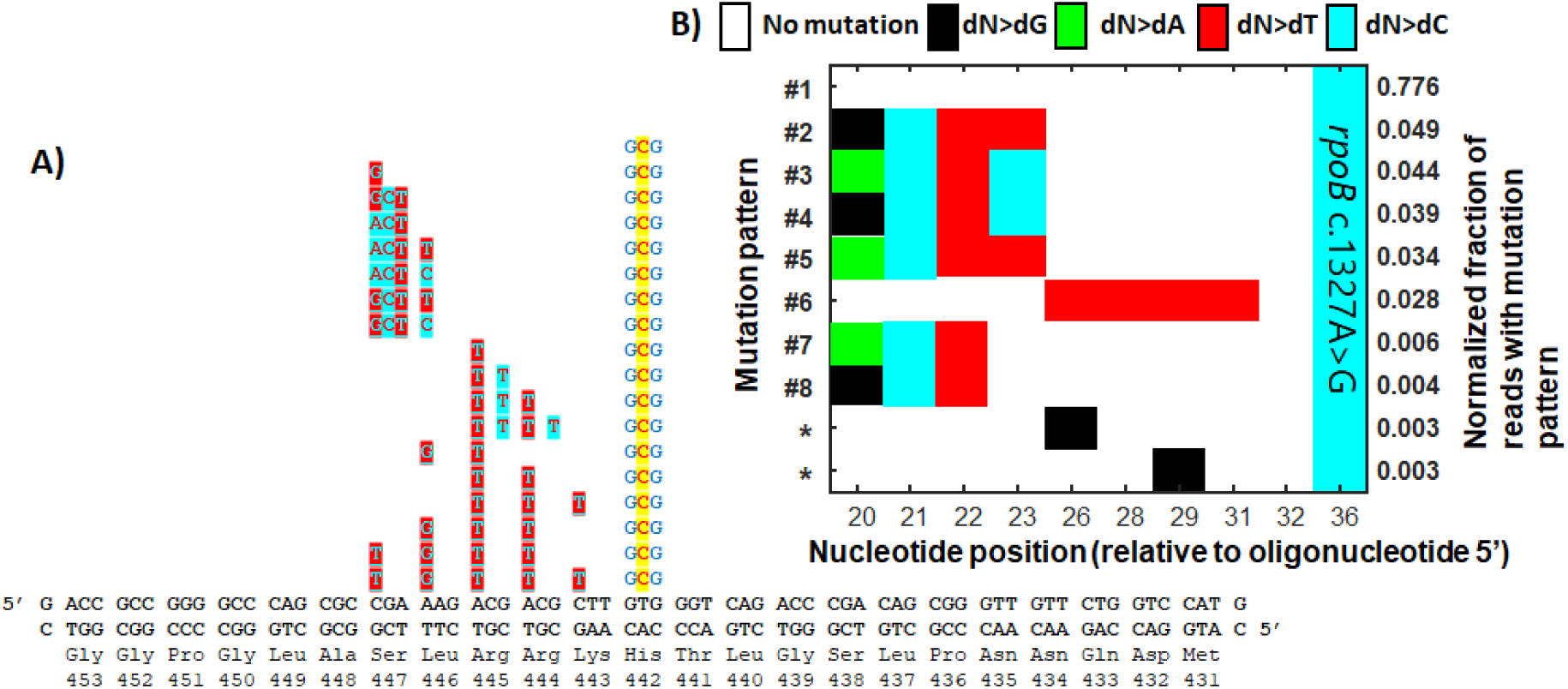
*Clustered mismatches are best able to allow NucS-active mismatches to evade repair during OR. A) Sequences of the pooled oligonucleotides evaluated for mutational patterns after rifampicin resistance. B) Mutational patterns (with relative frequency >0.1%) observed after sequencing. Oligonucleotides that introduce four consecutive mismatches, whether or not they are NucS-active or inactive, appear at the highest frequency, followed by 4 mismatches in across 6 nucleotides, followed by three consecutive mismatches. Patterns labelled with “*” represent unexpected mutational patterns observed that were not intentionally introduced by OR*.

## DISCUSSION

Our work, where mismatches are introduced specifically into the genome of *M. smegmatis*, provides evidence that NucS is involved in the direct repair of dT-dG mismatches *in vivo*, rather than, say, promoting the death of cells that are hyper-mutative for other reasons. Flanking NucS-inactive mismatches on the same oligonucleotide as a NucS-stimulating mismatch remain unrepaired and are converted to permanent genetic mutations if they are located >5 bp away from a NucS-stimulating mismatch, which suggests that the repair of the NucS-stimulating mismatch is confined within that small region. *In vitro*, NucS cleaves both strands two nucleotides 5’- of the mismatch (14,15,17), generating short sticky ends (Figure 7i), but we also observe NucS-inactive mismatches repaired “collaterally” if located three nucleotides away from NucS-activating mismatches. These findings are difficult to reconcile with the model of canonical RecA-dependent mycobacterial HR: they would not be observed for single-crossover HR (Figure 7), and a double-crossover tract would be required to observe such a mutational pattern—and that tract would have to be extraordinarily short; it would be short even in terms of fundamental limits of RecA filament assembly and homology recognition (45). The outcomes of repair more closely resemble repair by the mycobacterial NHEJ pathway (23,44,46), but with several differences: i) NHEJ tends to be highly mutagenic, including with insertions and deleted nucleotides at the site of repair, which we do not observe. Despite multiple possibilities for synonymous mutations at the sites of the dG-dT mismatches, we do not observe elevated mutation rate at that site to other nucleotides, but correct repair to dC. ii) There is no known mechanism for “strand discrimination” in the canonical NHEJ. In the absence of discriminating between 5’- overhangs caused by NucS, if one or the other 5’- overhang was being used to template repair, we might expect only 50% conversion, while we observe a near-total depletion of dT mutations at the sites of dT-dG mismatches corrected back to dC. These findings would all suggest that NucS is involved in a mechanistically-distinct MMR/DSB repair reaction that occurs within a short patch (Figure 7).

**Figure 7.**
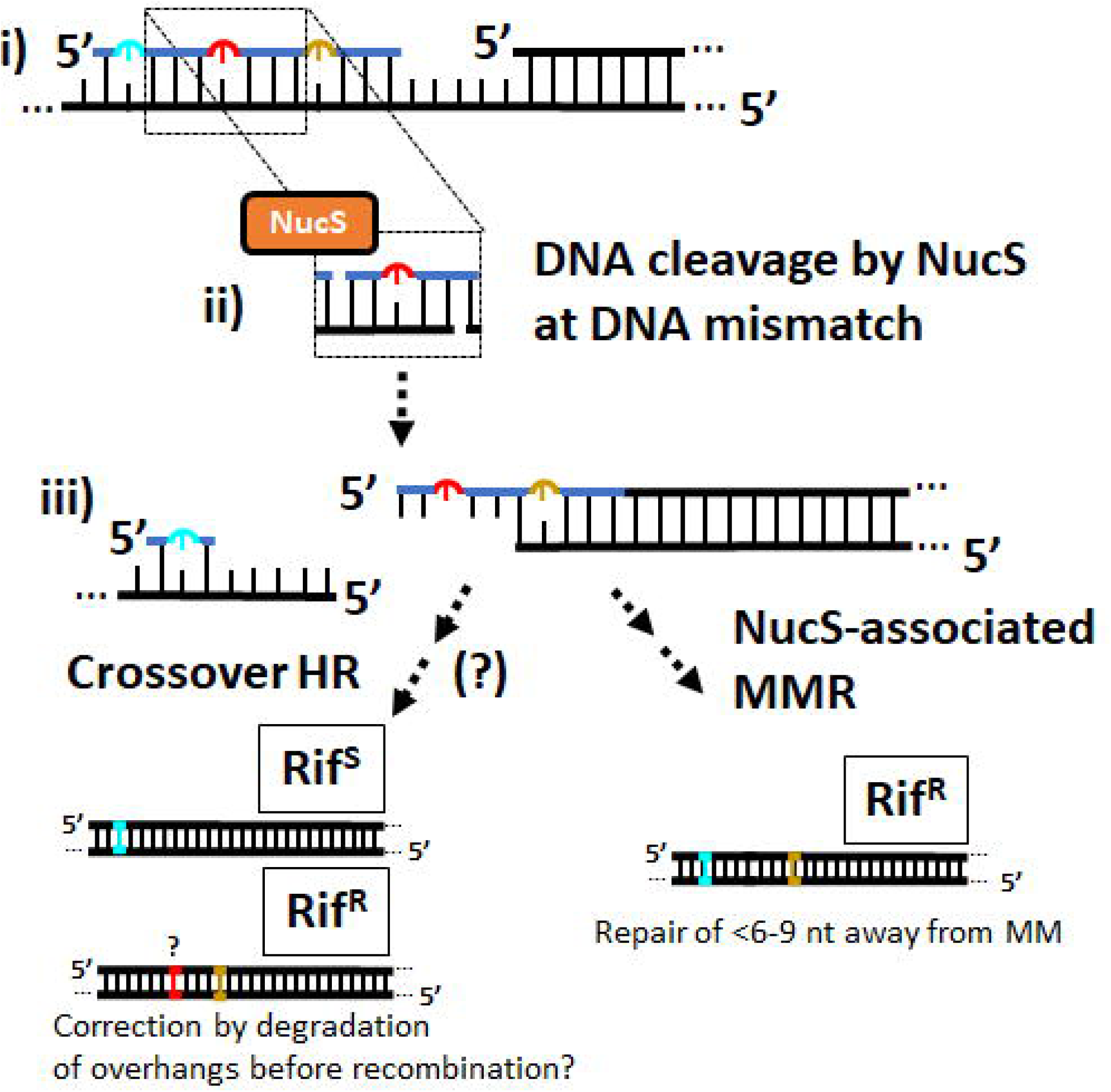
*Summary of experimental results. i) If during OR an oligonucleotide introduces a NucS-active dT-dG mismatch (red) to introduce a synonymous mutation is flanked at short distance by a NucS-inactive mutation to produce a synonymous mutation (blue) and NucS-inactive mutation that causes resistance to rifampicin (yellow), ii) NucS cleaves the dT-dG mismatch with two strand breaks located 2 nt 5’- of the mismatched dT and dG. iii) If repaired by homologous recombination (HR), we would not expect both of the mutations caused by the flanking mismatches to appear together with single-crossover repair events. However in a significant number of cases we find both flanking mutations with no corresponding mutation caused by the aggravating dT in between them*.

We note that our findings do not strictly eliminate the possibility of a NucS-mediated repair of mismatches that is mediated by HR or other DSB mechanisms that might also be present, and it is possible that degradation of the overhangs caused after NucS cleavage could result in complete repair using a sister chromatid. Mycobacterial helicase-nuclease AdnAB (47) that participates in HR has been shown to “nibble” a few 5’- nucleotides from sticky ends (48), and perhaps this is performed prior to HR after cleavage by NucS to remove both new and template sequences specifically at the site of mis-pairings. However, the observed 15% of correlated mutations flanking the sites of the dG-dT mismatch is likely a lower bound for the true amount of “short-patch” repair occurring: for example, the most prevalent mutation pattern (the *rpoB* c.1327A>G that causes rifampicin resistance alone) could be caused by repair by HR for some of the oligonucleotides we use (Figure 3D), but it also would be the expected product for all oligonucleotides that would cause flanking mis-pairs 3 nt away from the dG-dT mismatch if the same localized repair was occurring to those oligonucleotides. Likewise, those 50% of the oligonucleotides would introduce one flanking mis-pair 3 nt away from a dG-dT mismatch, which if repaired in a short patch might appear as a single mutation, which we also observe with relatively high frequencies.

A potential limitation of this study is that we were selecting for mutations in *rpoB*, an essential gene, and as such the repertoire of mutations that we would have the ability to observe by sequencing after repair—for example, frameshifts or large deletion—are less like to be observed as possible in a nonessential gene (23). This could potentially constrain the gene editing outcomes we can observe in this study, as those bacteria would not survive to be sequenced. However, that we do not see a significant difference in the rates of rifampicin resistance when oligonucleotides that introduce either NucS-active or NucS-inactive mis-pairs along with a NucS-inactive mis-pair that induces rifampicin resistance suggest that these types of repair outcomes are not common during repair. Furthermore, genetic mutation with the *rpoB* is an important cause of antibiotic resistance in *M. tuberculosis* and other mycobacterial pathogens (11). Having established OR as an experimental tool that can be used to probe mechanisms of NucS-mediated mutation avoidance in mycobacteria, and demonstrated ways to improve OR for mycobacterial genome engineering, it will be interesting to combine with other genetic tools (43,49-51) to further deconstruct the mechanism of mycobacterial MMR.

### MATERIAL AND METHODS

#### Preparation of *M. smegmatis* strains

*M. smegmatis* strain mc^2^155 (American Type Culture Collection) and *nucS*-knockout strain (prepared using ORBIT (43), a gift from Kenan Murphy of University of Massachusetts Chan Medical School) were prepared following the protocol of Ref. (30). Briefly, these strains were cultured in 7H9 broth (Millipore Sigma) containing Middlebrook ADC Growth Supplement, carbenicillin (50 ug/mL), cycloheximide (10 ug/mL), and 0.25% Tween 80 at 37°C until saturated (∼2 - 3 days). After 3 days of culture when the cells have reached OD600 0.8 − 1.0, 5 mL cultures were re-diluted into 50 mL 7H9 broth and transferred into large culture flask to grow overnight. Cells were pelleted by centrifuging at 3600g for 10 min at 4°C and the supernatant discarded, then the cells were washed with 1/2 vol (25 mL glycerol for 50 mL cell culture) 10% sterile glycerol by pipetting to disperse the cells. Washing the cells were performed in similar pattern with 1/4th, 1/8th, 1/10th volume of 10% sterile glycerol by centrifuging at 3600g for 10 min at 4°C with discarding the supernatant. The cells were resuspended in 1/25 vol (2 mL) 10% sterile glycerol to aliquot in 100 μL of cells in microcentrifuge tubes, and stored at −80 C for further experiment.

Electrocompetent cells were incubated for 10 min on ice, mixed with 50 ng of plasmid pJV62 (28) (pJV62 was a gift from Graham Hatfull (Addgene plasmid # 26910; http://n2t.net/addgene:26910; RRID:Addgene_26910)), and transferred to chilled 0.2 mm cuvette then electroporated using a BioRad pulse-controller (2.5 kV, 1000 Ω, 25 uF). The transformants were recovered in 7H9 broth then placed in a shaking incubator at 37°C shaker overnight. The cells were plated on 7H10 agar (Millipore Sigma) plates containing 50 ug/mL kanamycin and incubated at 37°C for 3-5 days until visible colonies appeared.

#### Oligonucleotide recombination

All experiments were performed in a manner similar to as described in Ref. (28) and in three replicates. 50 mL solutions of wild-type and *nucS* knock-out strains containing pJV62 plasmid were cultured in 7H9 broth with kanamycin for 3 days after which 2% (mass) acetamide to induce expression of Che9c gene 61 and allowed to grow for 24 hours. Bacteria were then made electrocompetent again following the above protocol prior to electroporation with 100 ng of oligonucleotides (either individually or pooled as described in the main text).

After transformation, the cultures were recovered overnight in 10 mL of 7H9 media and plated in 7H10 agar plate supplemented with 50 ug/mL rifampicin and incubated at 37°C until visible colonies appeared. For sequencing, instead of plating after the recovery step, 50 ug/mL rifampicin was added to each overnight culture directly, then incubated with shaking at 37°C.

After 3 days, 10 mL of cell cultures were transferred into tube to centrifuge at 3600 rpm for 15 min and the supernatant discarded. Cells were resuspended in 1 mL glucose-tris-EDTA (GTE) solution (25 mM Tris-HCl, pH 8.0, 10 mM EDTA, 50 mM glucose) then centrifuged for 10 min and the supernatant discarded. Cells were resuspended in 450 uL GTE solution with 50 uL of a 10 mg/mL lysozyme in 25 mM Tris-HCl and incubated at 37°C overnight. 100 uL 10% sodium dodecyl sulfate and 50 uL 10 mg/mL proteinase K were added to the cells and incubated at 55°C for 40 min, followed by the addition of 200 uL 5 M NaC, a gentle mixing, and the addition of 160 uL of a cetyltrimethylammonium bromide (CTAB) (10 g CTAB and 4.1 g NaCl in 90 mL dH2O.) was added and incubated at 65°C for 10 min. After 10 min, cells were washed with 1 mL chloroform:isoamyl alcohol (24:1) and centrifuged at 3600 rpm for 10 min. The supernatant was discarded and the sample washed again. The supernatant was discarded and 560 uL (0.7x by vol) isopropanol was added and mixed gently by inversion until the DNA has precipitated out of solution. This solution was incubated at room temperature for 5 minutes and centrifuged for 10 minutes. After discarding the supernatant, cells were washed with 1 mL 70% ethanol and centrifuged for 10 minutes. The supernatant was discarded and air-dried pellet for 15 min. The pellets were resuspended in 50 μL TE buffer and incubate at 37°C to dissolve pellet and concentrations were measured in Qubit fluorometer (Thermo Fisher). These extracted genomic DNA were stored at −20°C for further analysis

#### Sequencing and Analysis

For Sanger sequencing, PCR using the purified genomic DNA was performed with primers 5’-ACCGAAAAGGGCACCTTCAT-3’ and 5’-ACCGATCAGACCGATGTTGG-3’ to amplify the region of the *rpoB* gene of interest using OneTaq 2X Master Mix with Standard Buffer (NEB). Sanger sequencing was performed by Azenta Life Sciences (South Plainfield, NJ).

For next generation sequencing (NGS), PCR was performed with genomic DNA with primers containing partial illumina adapter sequences: 5’-ACACTCTTTCCCTACACGACGCTCTTCCGATCTCCGCAGACCCTGATCAACAT-3’ and 5’-GACTGGAGTTCAGACGTGTGCTCTTCCGATCTATCAGACCGATGTTGGGACC-3’. Sequencing was performed by Azenta Life Sciences using their Amplicon E-Z services, which guarantees >50,000 reads per sequencing run.

The lengths of the sequencing reads were measured using code written in-house for MATLAB (Mathworks; Natick, MA) by checking for the sequences immediately flanking those complementary to the oligonucleotides, 5’-GCCGGCGCGCTCACGGGACA-3’ and 5’- AACTGCGACAGCTGGCTGGT-3’. After confirming that nearly all of the repair products were full length (lacking indels), we determined mutation rates for all sequencing reads containing (i) the flanking sequences listed immediately above and (ii) the *rpoB* c.1327A>G mutation indicative of successful oligonucleotide incorporation by searching the sequencing using a regular expression (regexp) for the oligonucleotide sequence with wildcards ([A,T,C,G]) at sites where pooled oligonucleotides had differences from the wild-type sequence. The number of reads at the wildcard locations that differed from the wild-type sequence (mutations) were normalized to the number of reads with containing the *rpoB* c.1327A>G mutation. Reads and mutations were only considered valid if the forward and reverse reads across the entire oligonucleotide sequence were identical.

### SUPPORTING INFORMATION

Additional images of mycobacterial growth after OR, additional sequencing results, and lists of oligonucleotide sequences used are available in the Supporting Information

### DATA AVAILABILITY

Sequencing data and scripts for analysis are available upon request to the corresponding author and will be published online following peer review.

## ACKNOWLEDGEMENTS

We wish to acknowledge Kenan Murphy, PhD (UMass Chan Medical School) for supplying *M. smegmatis* strain *mc^2^155* (nucS<>attB) stain used for controls. pJV62 was a gift from Graham Hatfull (Addgene plasmid # 26910; http://n2t.net/addgene:26910; RRID:Addgene_26910).

We also thank Rachel Tinker-Kulberg and Tasmia Islam for technical assistance.

