## Supplementary Information for "Genome Editing Outcomes Reveal Mycobacterial NucS Participates in a Short-Patch Repair of DNA Mismatches"

**SUPPORTING INFORMATION**

**Figure S1.** All three replicates of the experiment described in Figure 1D.

**Figure S2.** Additional replicates of experiment described in Figure 2B (top) and 2C (bottom), respectively.

**Figure S3.** Histogram of read lengths between the sequences immediately outside the oligonucleotide sequences.

**Figure S4.** Variation of experiment performed in Figure 4, again showing that NucS-associated MMR collaterally repairs NucS-inactive mismatches within 6 nucleotides of a NucS-active mismatch but not outside 6 – 9 nt of the NucS-active mismatch.

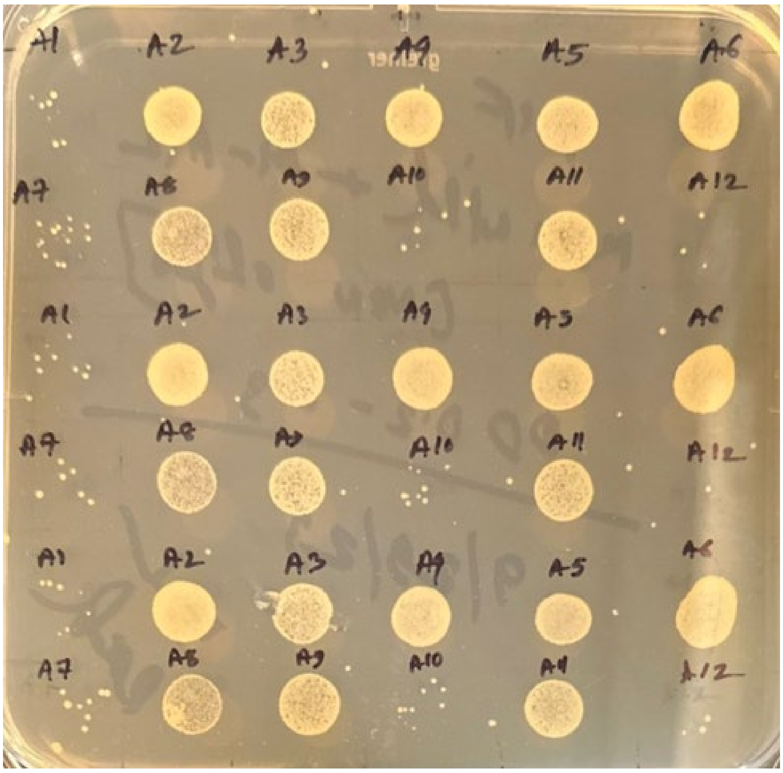

**Figure S1.** All three replicates of experiment described in Figure 1D.

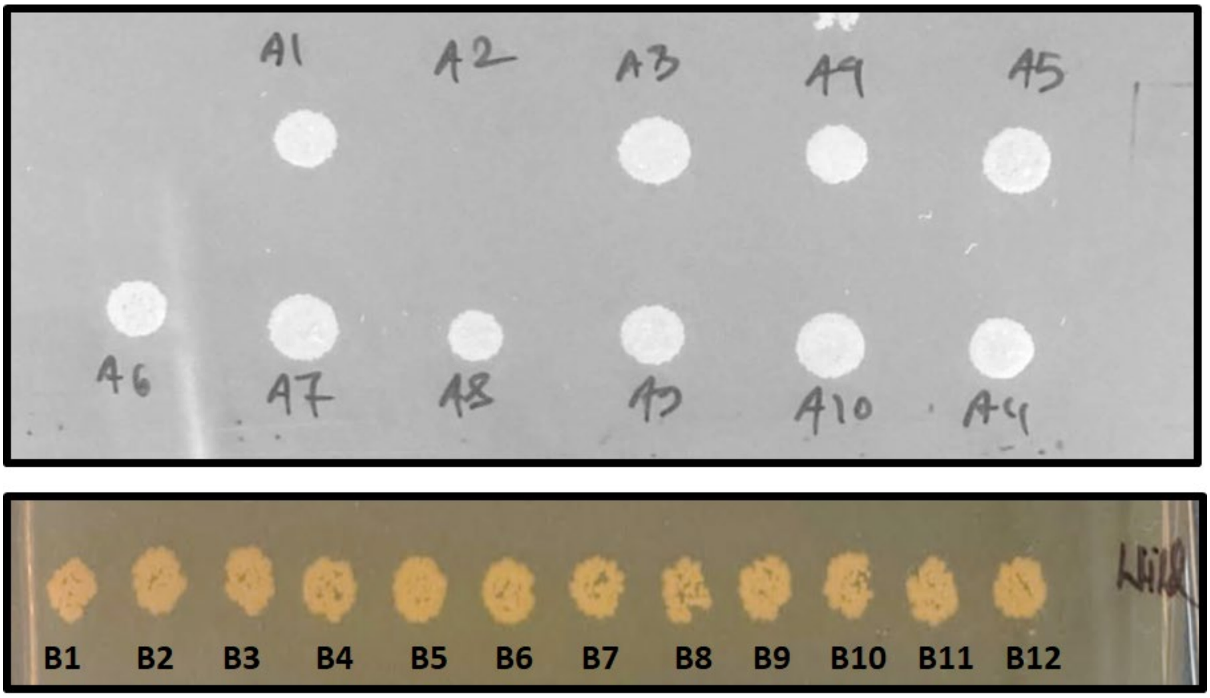


**Figure S2.** Additional replicates of experiment described in Figure 2B (top) and 2C (bottom), respectively.

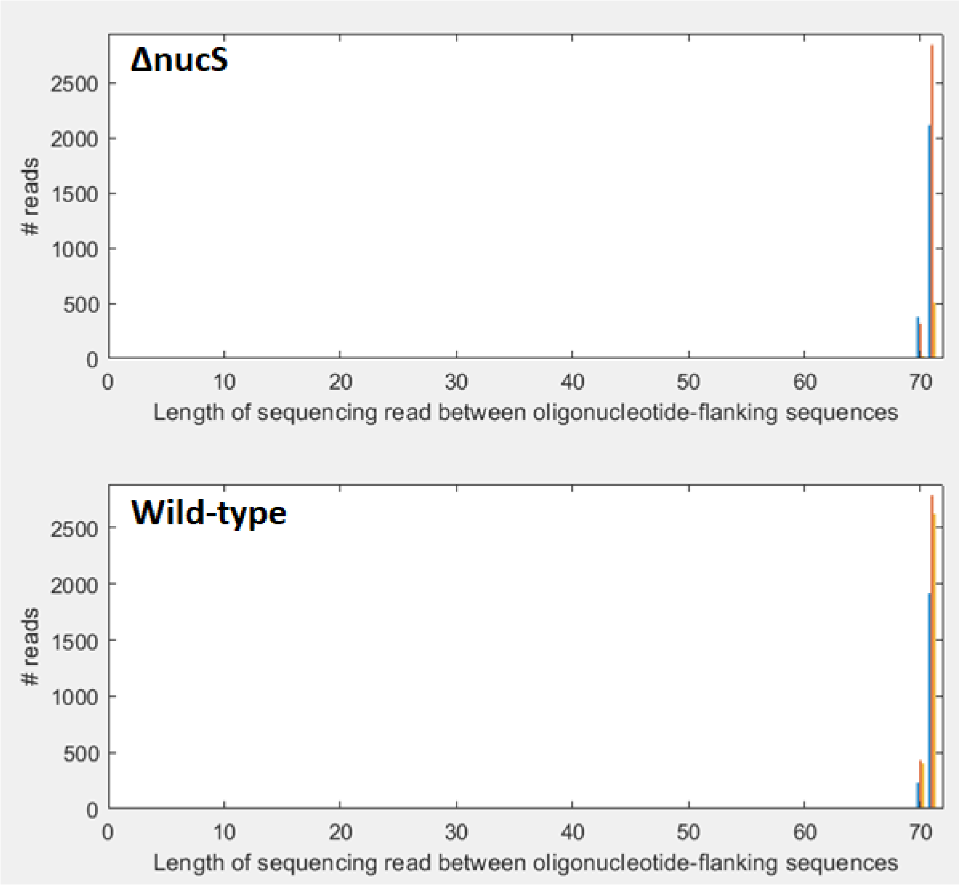


**Figure S3.** Histogram of read lengths between the sequences immediately outside the oligonucleotide sequences. Expected length 71 bp. Note there is a small amount of sequences that appear to be 70 bp: these appear in both NucS knockout and wild-type strains, and are very likely a sequencing artifact of a 'missing' nucleotide in a region of low complexity, away from the sites of introduced mutations. There is no evidence of insertions or deletions during repair.

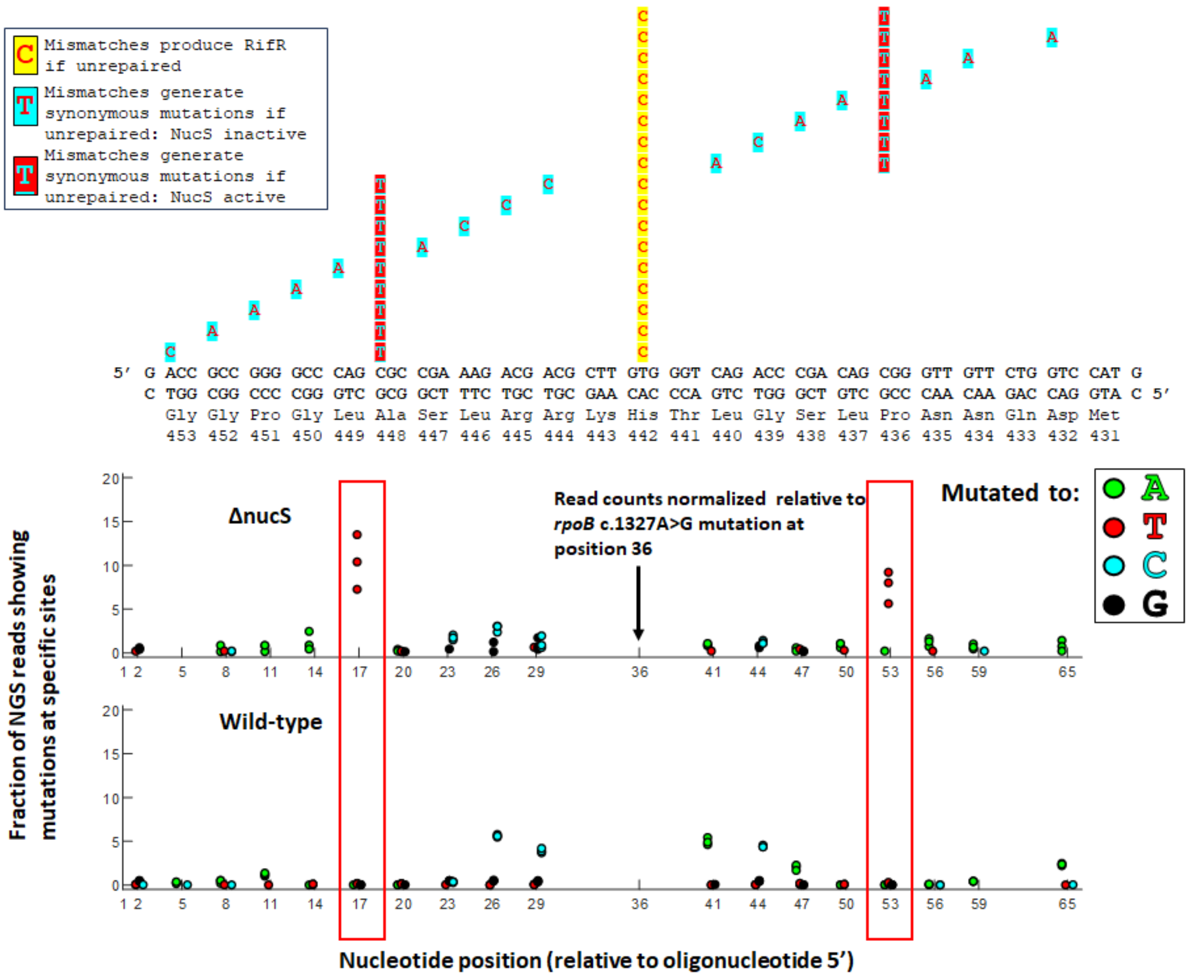


**Figure S4.** Variation of experiment performed in Figure 4, again showing that NucS-associated MMR collaterally repairs NucS-inactive mismatches within 6 nucleotides of a NucS-active mismatch but not outside 6 – 9 nt of the NucS-active mismatch**.** A) Pooled oligonucleotides (blue, see Figure 1 caption) that contain i) a dA-dC mismatch that should introduce a rifampicin resistant phenotype if unrepaired; ii) a NucS-active dT-dG mismatch located either 5′- or 3′- of (i) that would produce synonymous mutation if unrepaired; and iii) one NucS-inactive mismatches (e.g., dA-dC, dC-dC, dA-dA, dT-dC) that would produce synonymous mutations in rpoB if unrepaired, at various positions relative to (i) and (ii). B) Mutations generated by both NucS-active (boxed in red) and NucS-inactive mismatches within 3 nt of a NucS-active mismatch are significantly depleted in the NucS-active strain. Mutations generated by NucS-inactive mismatches > 6 nt away are largely unaffected, though there is a slight effect 9 nt 3′- of the NucS-active mis-pair. Note that the results presented show 3 biological replicates (if fewer than three dots are observed, it is because they are overlapping).
